# Subtle stratification confounds estimates of heritability from rare variants

**DOI:** 10.1101/048181

**Authors:** Gaurav Bhatia, Alexander Gusev, Po-Ru Loh, Hilary Finucane, Bjarni J. Vilhjálmsson, Stephan Ripke, Schizophrenia Working Group of the Psychiatric Genomics Consortium, Shaun Purcell, Eli Stahl, Mark Daly, Teresa R de Candia, Sang Hong Lee, Benjamin M Neale, Matthew C. Keller, Noah A. Zaitlen, Bogdan Pasaniuc, Nick Patterson, Jian Yang, Alkes L. Price

**Author notes:** Correspondence should be addressed to G.B. or A.L.P.

## Abstract

Genome-wide significant associations generally explain only a small proportion of the narrow-sense heritability of complex disease (*h*^2^). While considerably more heritability is explained by all genotyped SNPs (*h*_*g*_^2^), for most traits, much heritability remains missing (*h*_*g*_^2^ < *h*^2^). Rare variants, poorly tagged by genotyped SNPs, are a major potential source of the gap between *h*_*g*_^2^ and *h*^2^. Recent efforts to assess the contribution of both sequenced and imputed rare variants to phenotypes suggest that substantial heritability may lie in these variants. Here we analyze sequenced SNPs, imputed SNPs and haploSNPs— haplotype variants constructed from within a sample, without using a reference panel— and show that studies of heritability from these variants may be strongly confounded by subtle population stratification. For example, when meta-analyzing heritability estimates from 22 randomly ascertained case-control traits from the GERA cohort, we observe a statistically significant increase in heritability explained by imputed SNPs even after correcting for principal components (PCs) from genotyped (or imputed) SNPs. However, this increase is eliminated when correcting for stratification using PCs from a larger number of haploSNPs. We note that subtle stratification may also impact estimates of heritability from array SNPs, although we find that this is generally a less severe problem. Overall, our results suggest that estimating the heritability explained by rare variants for case-control traits requires exquisite control for population stratification, but current methods may not provide this level of control.

## Introduction

While genome-wide association studies (GWAS) have been extremely successful in identifying robust associations between single nucleotide polymorphisms (SNPs) and complex traits, the aggregate heritability explained by these loci (*h*_GWAS_^2^) is only a small fraction of the heritability estimated from related individuals (*h*^2^)^1^. This gap between *h*^*2*^ (ref. 2) and *h*_GWAS_^2^—termed the “missing heritability^3^”—is partially explained by causal variants that have not achieved genome-wide significance in GWAS of current sample sizes^4^. However, the heritability explained by all genotyped SNPs (*h*_*g*_^2^) typically leaves much of *h*^*2*^ unaccounted for and the pattern of *h*_GWAS_^2^ < *h*_*g*_^2^ < *h*^*2*^ is observed across a broad set of complex traits^1^. One possible explanation for the remaining missing heritability is that rare genetic variants—untyped by most genotyping arrays—explain a significant portion of the variance of studied traits^4,5^.

As sequencing data becomes more readily available, it will become possible to directly assess the heritability explained by rare variants. In fact, a recent analysis suggests that rare variants from previously associated regions of the genome^6^ may explain a substantial fraction of the heritability of prostate cancer. Even in the absence of sequence data it may be possible to assess the heritability explained by rare variants through imputation using a high coverage reference panel^7,8^. While accurately imputed SNPs do not typically explain more heritability than genotyped SNPs alone^9,10^, including low-accuracy imputed SNPs can explain significantly more of the heritability of quantitative traits^11^.

Despite the promise of analyses of rare variants, these analyses may be more affected by confounding than corresponding common variant analyses. In addition to sequencing artifact^12^ and imputation error^13^, analyses of rare variants may be particularly susceptible to subtle population stratification^14^ that cannot be corrected by standard techniques^15^. To assess this, we analyzed the estimates of genetic variance between the two control cohorts in the well-studied UK10K data-set^8^. Control cohort label should not be a heritable phenotype and any nonzero control-control heritability is considered a signal of uncorrected confounding^10,16^. In this analysis, we observed evidence of severe confounding for estimates from sequenced SNPs, imputed SNPs, and haploSNPs— haplotype variants constructed from within a sample, without using a reference panel. Indeed, estimates from all of these types of rare variants were often outside of the interpretable range (> 1). An even greater concern is that these estimates of control-control heritability from rare variants were largely unaffected by principal components (PCs) from array SNPs, sequenced SNPs, or imputed SNPs. Inclusion of haploSNP PCs produced statistically significant reductions in control-control heritability for all types of rare variants, but did not eliminate it entirely. This is consistent with previous analysis showing that analysis of haplotype structure can improve ancestry inference relative to SNPs alone^17^. We note that we also observed nonzero control-control heritability in estimates from array SNPs in the UK10K data; LD score regression^18^ correctly identified this control-control heritability as confounding, but is not designed for analyses of rare variants.

To assess whether confounding was also present in real case-control datasets we averaged estimates for 22 randomly ascertained complex diseases in the Genetic Epidemiology Research on Adult Health and Aging (GERA) cohort^19^. We observed significantly more heritability in imputed SNPs 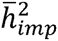 = 0.14 (s. e. 0.02) than in array SNPs alone (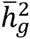 = 0.11 (s. e. 0.01); *P* = 0.01 for difference) when correcting for PCs computed from genotyped SNPs. Correcting for PCs computed from imputed SNPs did not affect these heritability estimates. However, inclusion of PCs computed from haploSNPs produced statistically significant reductions in both 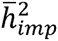 and 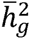, such that the difference between them was no longer statistically significant. We observe even larger evidence of confounding in ascertained case-control studies of schizophrenia (PGC2 cohort) and multiple sclerosis (WTCCC2 cohort). Overall, we conclude that estimating the heritability explained by rare variants for case-control traits requires exquisite control of population stratification, but current methods may not provide this level of control.

## Results

### UK10K control-control heritability estimates

Evidence of genetic differences between control cohorts has been previously used as a signal of confounding^10,16^ in estimates of heritability explained by array SNPs. We conducted control-control analyses using low-coverage (7x) sequencing data from the UK10K project^8^, which is comprised of two cohorts—TWINSUK and ALSPAC. The availability of sequence data allows us to assess control-control heritability using array SNPs that are present on standard genotyping arrays as well as rare variants that may be more susceptible to subtle population structure. As control cohort label is not expected to be a heritable phenotype, any significant control-control heritability is an indication of confounding. For all analyses we define the heritability explained by a set of markers analogously to previous work^20^, accounting for the possible presence of population structure (see Online Methods). All control-control estimates are on the observed scale and were produced using Haseman-Elston (HE) regression^21^. We note that PCGC regression^22^, a generalization of HE regression that produces estimates on the liability scale, is the main method used in our analyses of real case-control phenotypes in subsequent sections (see Online Methods and URLs).

### Heritability explained by array SNPs

We first estimated the heritability explained by array SNPs (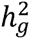) present on the Illumina Human660-Quad chip genotyping array (used by WTCCC2 ^23^). After restricting the UK10K data to these SNPs, we performed stringent quality control (see Online Methods), ultimately analyzing data from 3,565 individuals at 408k array SNPs. We used HE regression^21^ to estimate 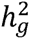 of the control-control phenotype (1 for the TWINSUK cohort; 0 for the ALSPAC cohort) at 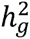 = 0.51 (s. e. 0.13), suggesting that subtle stratification results in substantial control-control heritability. Results did not change significantly when we included principal components (PCs) from array SNPs, which produced a similar estimate of 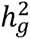 = 0.53 (s. e. 0.13) (see Table 1), or when we used the GCTA software package^24^ to estimate 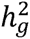 (see Table S1). We note that two sequencing centers contributed data to the UK10K project, but sequencing center was not correlated with cohort label, making control-control heritability due to batch effects unlikely. As array SNPs showed substantially less evidence of sequencing error^8^, we conclude that subtle population stratification rather than assay artifact is the most likely explanation of the observed confounding. We note that we also observed signals of control-control heritability from array SNPs in the well-studied WTCCC2^23^ data (see Table S2), even after excluding the possibility of assay artifact by restricting to genotypes that were concordant on separate genotyping arrays.

**Table 1.**
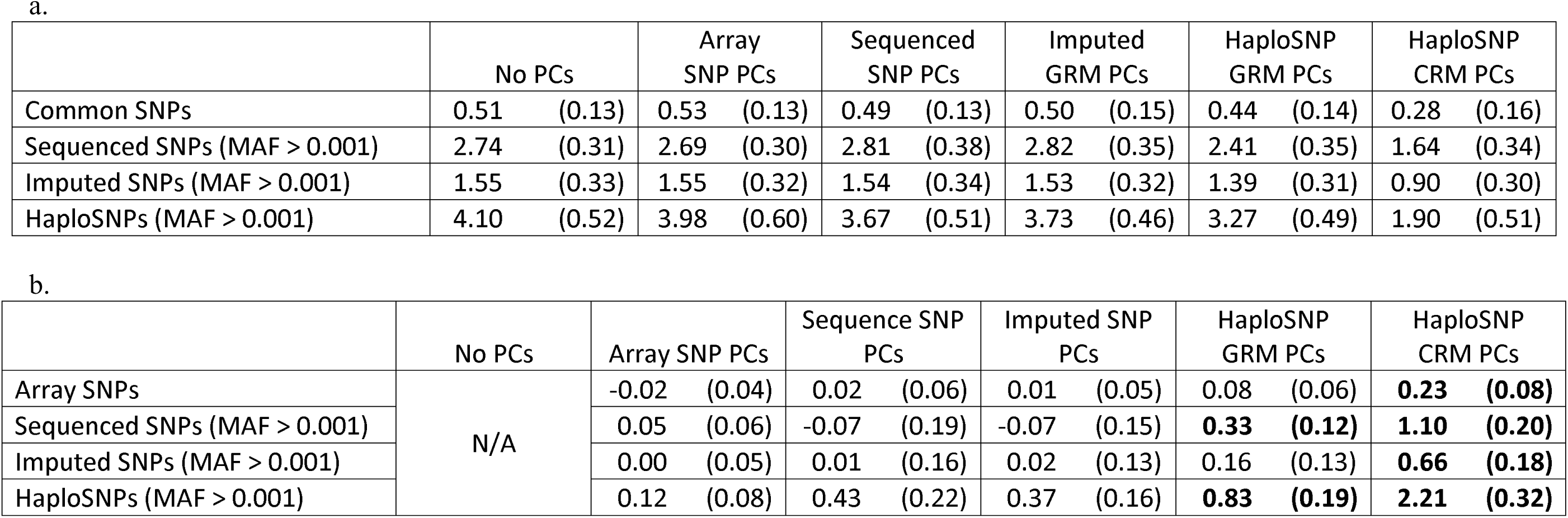
UK10K Control-Control Heritability Estimates with Different Corrections For Population Stratification. (a) We estimate control-control heritability explained by array SNPs and three types of rare variants: sequenced SNPs, imputed SNPs and haploSNPs. All types of variants show substantial evidence of confounding. Inclusion of PC covariates from array SNPs, sequenced SNPs, or imputed SNPs does not alter the estimate of heritability explained. While inclusion of haploSNP CRM PC covariates produces statistically significant drops in the heritability explained by all sets of variants, substantial evidence of confounding remains. (b) Given the small sample size of the UK10 data-set, standard errors on estimates are large. However, as analyses were based on the same individuals and, in some cases, the same set of variants, errors in the estimates were highly correlated. Thus, the standard error of the difference between two estimates was smaller than suggested by the nominal standard errors. We estimated the standard error of the difference (s.e.d.) for a pair of estimates using a jackknife over individuals (see Online Methods). The table above lists the difference and s.e.d. for an analysis with a set of PCs and the corresponding analysis with no covariates. For example, inclusion of haploSNP CRM PCs as covariates reduced the estimate from sequenced SNPs by 1.10 (s.e.d. 0.20) (from 2.74 (s.e. 0.31) with no covariates to 1.66 (s.e. 0.38) with haploSNP CRM PCs). Statistically significant reductions in control-control heritability are indicated in bold.

|  | No PCs |  | Array<br>SNP PCs |  | Sequenced<br>SNP PCs |  | Imputed<br>GRM PCs |  | HaploSNP<br>GRM PCs |  | HaploSNP<br>CRM PCs |  |
| --- | --- | --- | --- | --- | --- | --- | --- | --- | --- | --- | --- | --- |
| Common SNPs | 0.51 | (0.13) | 0.53 | (0.13) | 0.49 | (0.13) | 0.50 | (0.15) | 0.44 | (0.14) | 0.28 | (0.16) |
| Sequenced SNPs (MAF > 0.001) | 2.74 | (0.31) | 2.69 | (0.30) | 2.81 | (0.38) | 2.82 | (0.35) | 2.41 | (0.35) | 1.64 | (0.34) |
| Imputed SNPs (MAF > 0.001) | 1.55 | (0.33) | 1.55 | (0.32) | 1.54 | (0.34) | 1.53 | (0.32) | 1.39 | (0.31) | 0.90 | (0.30) |
| HaploSNPs (MAF > 0.001) | 4.10 | (0.52) | 3.98 | (0.60) | 3.67 | (0.51) | 3.73 | (0.46) | 3.27 | (0.49) | 1.90 | (0.51) |

**Table 1. UK10K Control-Control Heritability Estimates with Different Corrections For Population Stratification.**
|  | No PCs | Array SNP PCs |  | Sequence SNP<br>PCs |  | Imputed SNP<br>PCs |  | HaploSNP<br>GRM PCs |  | HaploSNP<br>CRM PCs |  |
| --- | --- | --- | --- | --- | --- | --- | --- | --- | --- | --- | --- |
| Array SNPs | N/A | -0.02 | (0.04) | 0.02 | (0.06) | 0.01 | (0.05) | 0.08 | (0.06) | **0.23** | **(0.08)** |
| Sequenced SNPs (MAF > 0.001) |  | 0.05 | (0.06) | -0.07 | (0.19) | -0.07 | (0.15) | **0.33** | **(0.12)** | **1.10** | **(0.20)** |
| Imputed SNPs (MAF > 0.001) |  | 0.00 | (0.05) | 0.01 | (0.16) | 0.02 | (0.13) | 0.16 | (0.13) | **0.66** | **(0.18)** |
| HaploSNPs (MAF > 0.001) |  | 0.12 | (0.08) | 0.43 | (0.22) | 0.37 | (0.16) | **0.83** | **(0.19)** | **2.21** | **(0.32)** |

We next sought to assess whether PCs computed from rare variants might be better able to correct for the confounding that we observed. We performed stringent quality control on UK10K sequence data, and computed PCs from 17.6M sequenced SNPs, excluding singletons (see Online Methods). We divided these SNPs into 7 MAF bins and computed 20 PCs from each bin, along with 20 PCs from array SNPs. Despite including these 160 PCs from sequenced and array SNPs, our estimate of 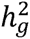 was essentially unchanged at 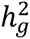 = 0.49(s.e. 0.13)(seeTable1). We repeated this analysis including 160 PCs from imputed and array SNPs, but did not observe any significant change in 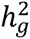 estimates (see Table 1).

Finally, we attempted to correct 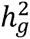 estimates for confounding using PCs computed from haploSNPs: variants defined using a new method that analyzes haplotype structure within the target sample, efficiently tagging unobserved rare variation without a reference panel (see Online Methods and URLs). Using computationally phased genotypes^25^ at array SNPs, we built haploSNPs from all pairs of phased chromosomes in the sample. To build haploSNPs, we began at a SNP at which the two chromosomes match, and extended the haploSNP one SNP at a time. The haploSNP was extended until a terminating mismatch—a mismatch that cannot be explained as a mutation on a shared background, or until a maximum length of 50kb is reached. Terminating mismatches were detected as violations of the 4-gamete test between the haploSNP being extended and the mismatch SNP. Using the array SNPs described above we constructed a set of 32.3M haploSNPs, excluding singletons. We observed that haploSNP GRMs were somewhat dominated by noise from outlier individuals, and inclusion of PCs from GRMs for each of 7 MAF bins of haploSNPs along with PCs from array SNPs did not substantially reduce heritability estimates (see Table 1). To reduce noise from outlier individuals, we computed correlation relationship matrices (CRMs) from haploSNP GRMs in each of 7 MAF bins, normalizing all entries by the appropriate diagonals (see Online Methods), and computed PCs from each haploSNP CRM. When we included 140 PCs from haploSNP CRMs along with 20 PCs from array SNPs as covariates, the estimate dropped to 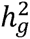 = 0.28 (s.e. 0.16) (see Table 1), which is no longer statistically significant. As analyses with and without haploSNP CRM PCs were highly correlated (i.e. based on the same individuals and variants), the reduction in 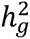 (0.23 (s.e. 0.08)) was statistically significant. We note that inclusion of PCs from sequenced or imputed SNP CRMs did not reduce estimates. Increasing the maximum length of haploSNPs allowed us to compute PCs that were significantly better at controlling for confounding (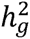 = 0.06 (s. e. 0.15); see Table S3), however, due to computational considerations in larger data sets, we report results for haploSNPs shorter than 50kb. Using phenotypes simulated without stratification from sequence SNPs, we confirmed that the observed drop in heritability estimates could not be explained by biases introduced by inclusion of a large number of PCs (see Table S4).

We also sought to assess whether LD score regression^18^ could correctly identify control-control heritability explained by array SNPs as confounding. We computed association statistics for array SNPs to the control-control phenotype without covariates. These statistics were substantially inflated (mean *χ*^2^ = 1.032) consistent with expectation given the sample size (*N* = 3,565), 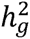 = 0.51 (s.e. 0.13), and assuming an effective number of SNPs of 60k^26^, since 1 + 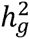 *N*/*M*_*EFF*_ = 1.030 (ref. 27). LD score regression assigned the bulk of the inflation in *χ*^2^ statistics to the intercept (1.031 (s.e. 0.007)), consistent with confounding. This suggests LD score regression may be able to detect confounding in heritability estimates from array SNPs even if inclusion of PCs from array SNPs and rare variants are insufficient to correct for it. (We note that attenuation bias can also produce low levels of inflation in the LD score regression intercept^18^.)

Another published strategy for assessing confounding in heritability estimates is comparison of genome-wide heritability estimates to the sum of estimates for each chromosome^28^. This relies on the idea that population stratification will be captured redundantly by SNPs on different chromosomes, inflating the sum of per-chromosome estimates relative to the genome-wide estimate. When we applied this strategy to per-chromosome estimates of control-control heritability, we observed no elevation of the sum of per-chromosome estimates (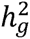 = 0.52 (s.e. 0.13)) relative to the genome-wide estimates (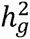 = 0.51 (s. e. 0.13)) (see Table S5). These results are consistent with simulations showing that this method may not always be well powered to detect subtle stratification in genome-wide estimates of heritability (see Table S6).

### Heritability explained by sequenced and imputed SNPs

Given the confounding of heritability estimates from array SNPs, we sought to assess whether heritability estimates from rare variants were also affected. We performed stringent quality control on UK10K sequence data (see Online Methods), and estimated heritability explained by sequenced SNPs with MAF > 0.001 (estimates from rarer variants were unstable; see Table S7). We split 11.7M sequenced variants (MAF > 0.001) into 5 MAF bins and used MAF-stratified HE regression to estimate control-control 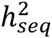 = 2.74 (s. e. 0.31)—an extremely confounded estimate outside the interpretable range from 0 to 1 (see Table 1 and Online Methods). Including 20 PCs from array SNPs (see above), or 160 PCs from sequenced and array SNPs did not change estimates substantially (see Table 1). However, when we included 140 PCs from haploSNP CRMs along with 20 PCs from array SNPs (see above) we observed a substantial reduction to 924 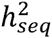 = 1.64 (s. e. 0.34). Estimates were consistent with those obtained from GCTA^24^ (see Table S1). Our results suggest that these PCs capture subtle population structure more effectively than PCs from other variants, though severe confounding remains. We note that CRM PCs from a larger number of longer haploSNPs further improved our ability to control for confounding (see Table S3), but were not able to remove the effects of confounding entirely.

To exclude the possibility that cohort-specific sequencing error at rare variants was the main source of confounding, we analyzed rare variants imputed from array SNPs. Specifically, we restricted to the set of 408k array SNPs described above, computationally phased these SNPs and imputed 13.0M rare variants (MAF > 0.001) using the Impute2 software package^29^ with 1000 Genomes data^30^ as a reference panel (see Online Methods). As previously suggested, we did not impose any imputation quality threshold^11^, maximizing our ability to tag untyped SNPs. While heritability estimates from imputed SNPs were somewhat lower, they remained severely confounded at 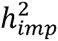 = 1.55 (s. e. 0.33) and were not reduced by inclusion of PCs from array SNPs, sequenced SNPs or imputed SNPs (see Table 1). Again, inclusion of PCs from haploSNP CRMs (along with array SNPs) produced a statistically significant reduction in 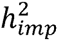 = 0.90 (s. e. 0.30), but did not eliminate confounding.

### Heritability explained by haploSNPs

Finally, to exclude the possibility that a combination of sequencing and imputation error were driving control-control heritability estimates from these variants, we analyzed the set of haploSNPs for the purposes of heritability estimation. We restricted to the set of 26.5M haploSNPs with MAF > 0.001, and estimated heritability explained by these variants using MAF-stratified HE regression as above. Our simulations without population stratification indicate that haploSNPs can effectively tag unobserved sequenced SNPs in the UK10K data (see Supplementary Note).

Of all types of rare variants, haploSNPs showed the strongest evidence of confounding: 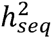 = 4.10 (s. e. 0.52); this confounding produced an inflation in the sum of per-chromosome estimates of heritability relative to the genome-wide estimate (see Table S5). Consistent with previous analyses, PCs from haploSNP CRMs produced the largest reduction.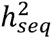 = 1.10 (s. e. 0.51); The fact that computationally inferred rare variants, unaffected by sequencing or imputation error, showed extreme confounding further implicates subtle population stratification rather than assay artifact as the most likely source of confounding. Indeed, previous work has noted that rare variants are subject to more local patterns of stratification^14^, which may be difficult to correct using standard methods.

### Recommendations

We recommend that estimates of heritability from array SNPs should be checked for confounding using LD score regression^18^. While there are alternate causes of inflation in the LD score regression intercept (e.g. attenuation bias or LD-dependent genetic architectures), a lack of observed inflation indicates that confounding due to population stratification is unlikely to be a major concern. If LD score regression shows evidence of confounding and/or there are other reasons for concern about stratification, PCs from haploSNP CRMs should be included as covariates, in addition to PCs from array SNPs. If a reduction is observed, CRM PCs from a larger number of longer haploSNPs can be investigated.

For heritability estimates from rare variants—sequenced SNPs, imputed SNPs, or haploSNPs—PCs computed from sequenced or imputed SNPs are unlikely to impact estimates. While PCs from haploSNP CRMs are more likely to detect subtle stratification, they are unlikely to correct for confounding completely. Thus, we recommend including PCs from haploSNP CRMs as covariates, but caution that any significant drop in estimates after inclusion of these PCs should be considered as evidence of uncorrected confounding rather than the resolution of this confounding. Overall, we believe that it is currently not possible to ensure that estimates of heritability from rare variants are robust to subtle population stratification.

## Analysis of 22 complex diseases (GERA cohort)

### Heritability explained by array SNPs

We next sought to assess the extent to which the subtle stratification observed in the UK10K control-control analyses might confound estimates of heritability from array SNPs in real case-control phenotypes. We focused on 22 complex diseases in 47,360 randomly ascertained samples typed at 289k SNPs in the GERA cohort^19^ (see Online Methods). Estimates were obtained for each disease using PCGC regression^22^, assuming that the prevalence of each disease matched the case fractions in the GERA cohort. Given the relatively low values of 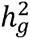 for any one trait^31^, we computed an average of liability-scale 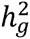 estimates across 22 traits (all averages across traits are inverse-variance weighted). We obtained an estimate of 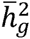 = 0.11 (s. e. 0.01) after including PCs from array SNPs (see Figure 1 and Table S8). We first sought to assess whether residual confounding could be impacting these estimates by applying LD score regression to association statistics (mean *χ*^2^ across all studies = 1.027) computed for each disease. LD score regression produced an average intercept of 1.005 (s.e. 0.002), suggesting that confounding is limited. This is consistent with our intuition that randomly ascertained studies are less susceptible to effects of stratification and assay artifact than ascertained case-control studies (see below).

**Figure 1.**
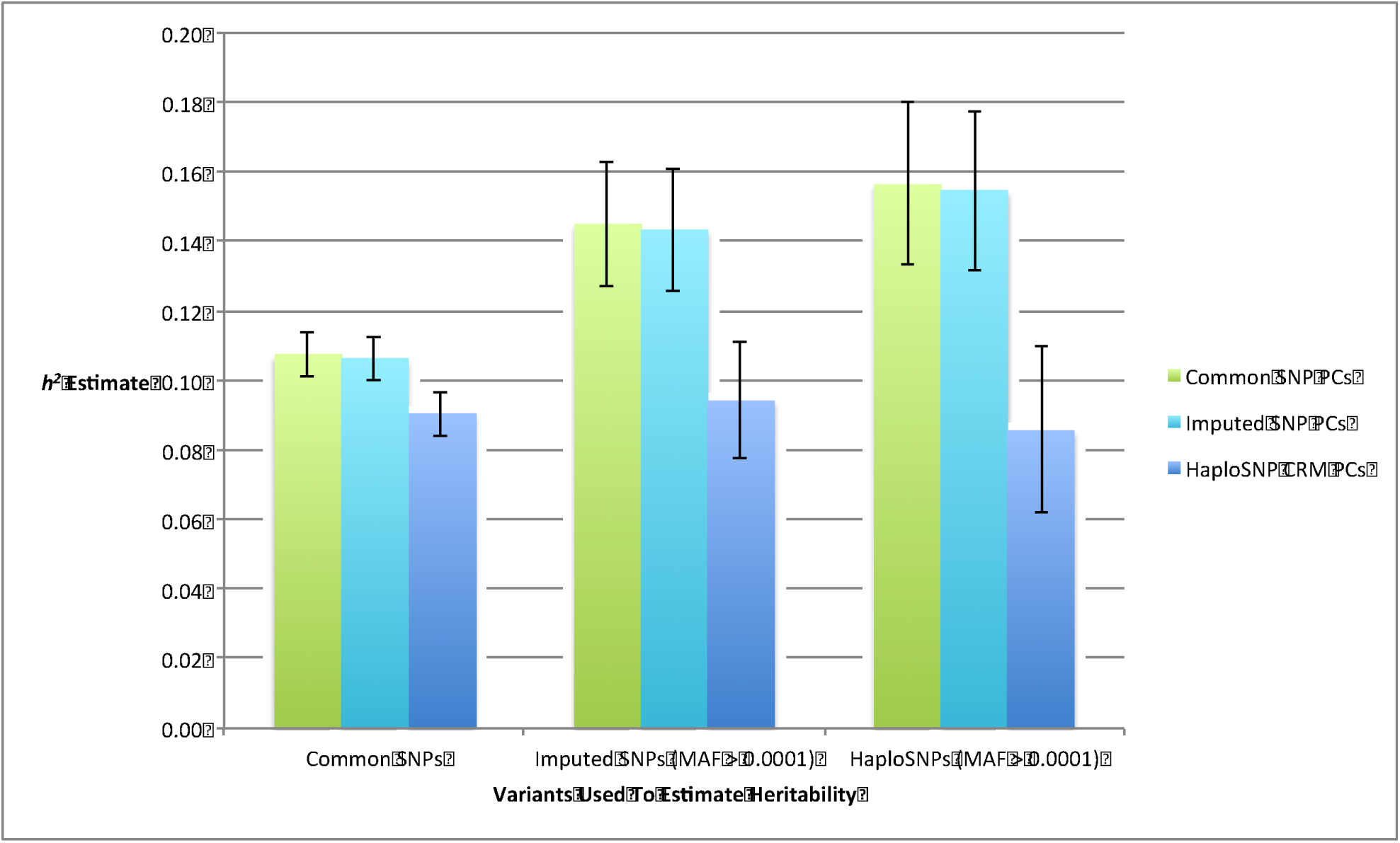
Estimates of heritability explained by common SNPs, imputed SNPs and haploSNPs, averaged across 22 traits in the GERA cohort. We estimated heritability explained by array SNPs, imputed SNPs and haploSNPs in the GERA data set (all estimates on the liability scale). Estimates are averaged using an inverse-variance weighted average of estimates for 22 traits (see Table S8 for individual trait estimates) and standard errors are displayed on the figure. After including array SNP PC covariates, both imputed and haploSNPs explain significantly more heritability than array SNPs alone. However, inclusion of haploSNP PCs eliminates all signals of rare variant heritability. This suggests that this data-set contains no signal of rare variant heritability beyond what is due to stratification.

We computationally phased these genotyped SNPs^25^, constructed 39.3M haploSNPs (MAF > 10^-4^) and computed 20 CRM PCs from each of 7 haploSNP MAF ranges (see Online Methods). Including PCs from haploSNP CRMs (along with PCs from array SNPs) reduced the estimate to 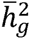 = 0.09 (s.e. 0.01), a difference of 0.015. Though quantitatively small, this difference was statistically significant (standard error of the difference (s.e.d.) 0.002 across 22 traits; see Table S9 and S10).

### Heritability explained by imputed SNPs

We sought to assess whether imputed SNPs or haploSNPs could explain significantly more heritability than array SNPs alone. We began by performing imputation using the Impute2 software package^29^ with a reference panel from the 1000 Genomes Project^30^ (see Online Methods). As previously described^11^, we did not impose any imputation quality filter, maximizing our ability to tag untyped causal variants. As this is a randomly ascertained study, assay artifacts are unlikely to be correlated with phenotypes and observed increases are likely to be due to polygenic signal or subtle population stratification. We estimated 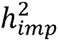 using MAF-stratified PCGC regression^22^ (see Online Methods). Averaged across 22 traits, we observe 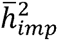 = 0.14 (s.e. 0.02) after correcting for stratification with PCs from array SNPs, a statistically significant increase of 0.038 over 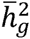 (s.e.d. 0.016; *P* = 0.02). Correcting for stratification with CRM PCs from haploSNPs (along with PCs from array SNPs) reduced our estimate to 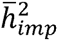 = 0.09 (s. e. 0.02) (see Figure 1 and Table S8), no longer showing any increase over the corresponding estimate of 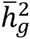(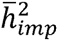 – 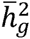 = 0.003; s.e.d. 0.016). We note that estimates of 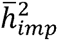 with and without PCs from haploSNP CRMs are strongly correlated as they are made using the same set of individuals and variants. Thus, the reduction in 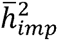 after inclusion of these PCs has a low s.e.d. (0.005, smaller than the s.e.d. of 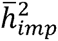 – 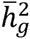) and is highly statistically significant (see Table S9). However, inclusion of PCs from imputed SNPs did not alter estimates (see Figure 1). Thus, an analysis utilizing current methods to correct for population stratification would have incorrectly concluded that imputed SNPs explained significantly more heritability than array SNPs alone.

### Heritability explained by haploSNPs

We estimated the heritability explained by the set of haploSNPs described above. Averaged across 22 traits we estimated 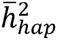 = 0.16 (s.e. 0.02) after correcting for PCs from array SNPs and 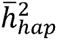 = 0.09 (s. e. 0.02) after correcting for PCs from haploSNP CRMs (along with PCs from arrays SNPs). Again, after correcting for PCs from haploSNP CRMs the difference 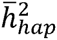 – 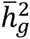 (average:-0.004; s.e.d. 0.023) was no longer statistically significant. Additionally, correcting for PCs from imputed SNP did not alter estimates substantially (see Figure 1 and Table S8), confirming that estimates of heritability explained by rare variants remain confounded even after applying standard methods to correct for population stratification.

Overall, our results suggest that confounding in estimates of heritability explained by array SNPs in randomly ascertained cohorts is limited, but that subtle stratification can produce spurious signals of heritability explained by rare variants. Additionally, confounding in estimates of heritability explained by rare variants cannot be appropriately corrected for through the inclusion of PCs from array SNPs or imputed SNPs as covariates in the analysis. After correction for PCs from haploSNP CRMs, we do not observe any signal of heritability from rare variants in this data set, but we caution that if such a signal were observed its robustness to subtle stratification would be unclear. Analysis of schizophrenia (PGC2 data)

### Heritability explained by array SNPs

We next sought to investigate whether issues related to case-control ascertainment— assay artifact correlated with phenotype or induced population stratification— would produce stronger confounding in estimates of heritability explained by array SNPs. We analyzed the heritability explained by array SNPs in PGC2-SCZ data^32^. We meta-analyzed estimates for each of ten cohorts of European ancestry with >1,000 individuals, for a total of >35,000 individuals (all averages across cohorts are inverse-variance weighted). We applied stringent quality control to genotyped SNPs, obtaining an average of 461k genotyped SNPs in each cohort (see Online Methods and Table S10). We estimated 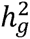 using PCGC regression^22^ (see Online Methods) using a disease prevalence of 1%^31^. For two studies of treatment resistant schizophrenia a disease prevalence of 0.3% was used. (We note that the choice of disease prevalence affects the absolute estimates, but not their relative values.) We meta-analyzed cohort-specific estimates (see Table 2), producing an average of 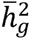 = 0.38 (s.e. 0.02) after includingarray SNP PCs as covariates, significantly larger than a previously published estimate^31^ 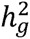 = 0.27 (s. e. 0.007)) computed from PGC2-SCZ samples (see Table 2).

**Table 2.** Estimates of the heritability of schizophrenia explained by array SNPs and haploSNPs in PGC2 cohorts. We analyze the heritability of schizophrenia explained by array SNPs and haploSNPs with MAF > 0.001 (all estimates on the liability scale). Estimates are meta-analyzed using an inverse-variance weighted average of cohort-specific estimates. All cohorts analyzed contained a minimum of 1000 samples (see Supplementary Material of ref. 32). Relative to array SNP PCs, inclusion of haploSNP PCs produces significant reductions in heritability estimated from haploSNPs and array SNPs. The meta-analyzed estimate made with^9,31^ array SNP PC covariates is substantially higher than previously reported estimates from mega-analysis of overlapping data-sets^9,31^. Our results suggest that cohort-specific population stratification, diluted in large mega-analyses may explain some of this difference. As before, we observe a large drop in 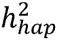 estimates after including haploSNP CRM PC covariates, but cannot conclude that residual confounding does not continue to inflate estimates. We also give the *P* value for heterogeneity of estimates across cohorts based on Cochran’s Q statistic^50^ and note that estimates of 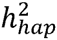 no longer show statistically significant evidence of heterogeneity after inclusion of PCs from haploSNP CRMs as covariates

| Cohort | Heritability Explained by Array SNPs |  |  |  | Heritability Explained by haploSNPs |  |  |  |
| --- | --- | --- | --- | --- | --- | --- | --- | --- |
|  | Array SNP PCs |  | HaploSNP CRM PCs |  | Array SNP PCs |  | HaploSNP CRM PCs |  |
| ajsz | 0.52 | (0.13) | 0.28 | (0.12) | 0.98 | (0.27) | 0.45 | (0.28) |
| boco | 0.43 | (0.08) | 0.26 | (0.08) | 2.14 | (0.30) | 1.04 | (0.28) |
| clm2 | 0.35 | (0.03) | 0.27 | (0.03) | 0.88 | (0.11) | 0.45 | (0.09) |
| clo3 | 0.46 | (0.06) | 0.29 | (0.07) | 1.64 | (0.26) | 0.67 | (0.21) |
| gras | 0.54 | (0.15) | 0.29 | (0.16) | 2.37 | (0.48) | 1.09 | (0.60) |
| irwt | 0.66 | (0.13) | 0.24 | (0.14) | 3.82 | (0.51) | 0.68 | (0.52) |
| mgs2 | 0.34 | (0.06) | 0.29 | (0.07) | 0.68 | (0.20) | 0.60 | (0.19) |
| s234 | 0.27 | (0.08) | 0.21 | (0.07) | 1.31 | (0.30) | 0.75 | (0.25) |
| swe5 | 0.38 | (0.07) | 0.31 | (0.08) | 0.41 | (0.23) | 0.29 | (0.25) |
| swe6 | 0.21 | (0.15) | 0.02 | (0.17) | 0.06 | (0.40) | -0.20 | (0.49) |
| **Average** | 0.38 | (0.02) | 0.27 | (0.02) | 1.03 | (0.07) | 0.52 | (0.06) |
| **Heterogeneity $P$** | 0.14 |  | 0.95 |  | **7.87E-12** |  | 0.43 |  |

To assess the degree of residual confounding, we computed association statistics including array SNP PCs as covariates and ran LD score regression^18^. Summary statistics were significantly inflated (mean *χ*^2^ across all studies = 1.048), and LD score regression estimated an average intercept of 1.014 (s.e. 0.003) indicating that residual confounding may be a concern (see Table S10), or based on a previously published intercept^18^ computed from PGC2-SCZ samples^32^. We note that the previous intercept was computed using association statistics from the combined set of 34,241 cases and 45,604 controls in which inflation due to cohort-specific population stratification, as observed in our analysis, would be diluted.

We next assessed whether PCs from haploSNP CRMs could better correct for this confounding by computationally phasing these genotypes^25^ and constructing an average of 32.8M haploSNPs (MAF > 10^-4^) in each cohort. We note that no cohort had more than 10,000 individuals, thus, we restricted our analysis to non-singleton haploSNPs. We computed 20 PCs from haploSNP CRMs for each of 7 MAF ranges (see Online Methods), and included these as covariates (along with PCs from array SNPs) in estimating 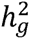 (see Table 2). Inclusion of these PCs significantly reduced the meta-analyzed estimate by approximately 30% 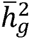 = 0.27 (s.e. 0.02)), now consistent with the estimate from ref. 31. This further confirms that PCs from haploSNP CRMs can be useful in correcting for confounding in estimates of heritability explained by array SNPs.

### Heritability explained by haploSNPs

We estimated the heritability explained by the set of haploSNPs described above. We focused solely on haploSNPs with MAF > 0.001, given the small sample sizes of our individual cohorts (see Table S10). Correcting for PCs from array SNPs, we obtained a meta-analyzed estimate of 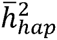 = 1.03 (s. e. 0.07) that showed a high degree of heterogeneity (*P* = 7.9x10^-12^) across cohorts (see Table 2). After including PCs from haploSNP CRMs as covariates, the meta-analyzed estimate dropped to 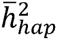 = 0.52 (s. e. 0.06) and estimates no longer showed significant of heterogeneity (*P* = 0.43 n.s.) (see Table 2 and Online Methods).

Our results suggest that estimates of heritability from rare haploSNPs are confounded by uncorrected population stratification. While including PCs from haploSNP CRMs reduced these estimates substantially, we cannot be certain that they correct for this confounding entirely and believe that estimates of heritability explained by rare haploSNPs should be viewed with caution. Notably, confounding is substantially more severe in estimates from this ascertained case-control data than in the randomly ascertained GERA cohort (see above). We note that a strategy of assessing cross-cohort heritability explained by rare variants may be a promising approach to separating the effects of cohort-specific confounding and true polygenicity. However, subtle stratification drives differentiation between cohorts, particularly at rare variants. In a setting with stratification, cross-cohort analyses may actually estimate heritability that is shared across populations and produce overly conservative estimates as a result (see Supplementary Note and Table S11).

## Analysis of multiple sclerosis (WTCCC2 data)

### Heritability explained by array SNPs

We analyzed the genome-wide heritability explained by array SNPs in the WTCCC2-MS data set^23,26^. We used extremely stringent quality-control filters to avoid inflation due to assay artifacts^10,16^, and all analyses excluded chromosome 6 (see Online Methods). We note that while there is a large ancestry mismatch between WTCCC2 MS cases and controls as a consequence of the set of samples that are publicly available^26^, estimates of 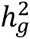 have been previously obtained in this data^10,26^ by including PCs from array SNPs as covariates, and by analyzing an ancestry-matched subset of the data^10^. Using PCGC regression^22^, we estimated 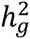 = 0.27 (s. e. 0.02), consistent with prior estimates^26,33^. All estimates reported are on the liability scale assuming a disease prevalence of 0.1%^33^.

To assess the degree of residual confounding at array SNPs, we computed association statistics including PCs from array SNPs as covariates and ran LD score regression^18^. Association statistics were substantially inflated overall (mean *χ*^2^ = 1.15), and LD Score regression assigned a large fraction of this inflation to the intercept term (1.06 (s.e. 0.009)). This suggests that substantial uncorrected population stratification confounds our estimate of 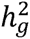, despite the inclusion of PCs from array SNPs as covariates. To assess whether inclusion of PCs computed from rare haploSNPs could correct for this stratification we computationally phased this set of genotyped SNPs^25^, built a set of 53.0M haploSNPs (MAF > 10^-4^), and computed 20 PCs from haploSNPs in each of 7 MAF ranges (see Online Methods). After including PCs from haploSNP CRMs (along with PCs from array SNPs) we estimated 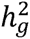 = 0.17 (s. e. 0.02) (see Figure 2; Online Methods). While we cannot exclude the possibility that some uncorrected stratification could confound this estimate, we believe that 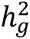 = 0.17 (s. e. 0.02) is a more accurate estimate of the 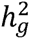 of multiple sclerosis in this data set than the larger values reported previously^10,26^.

**Figure 2.**
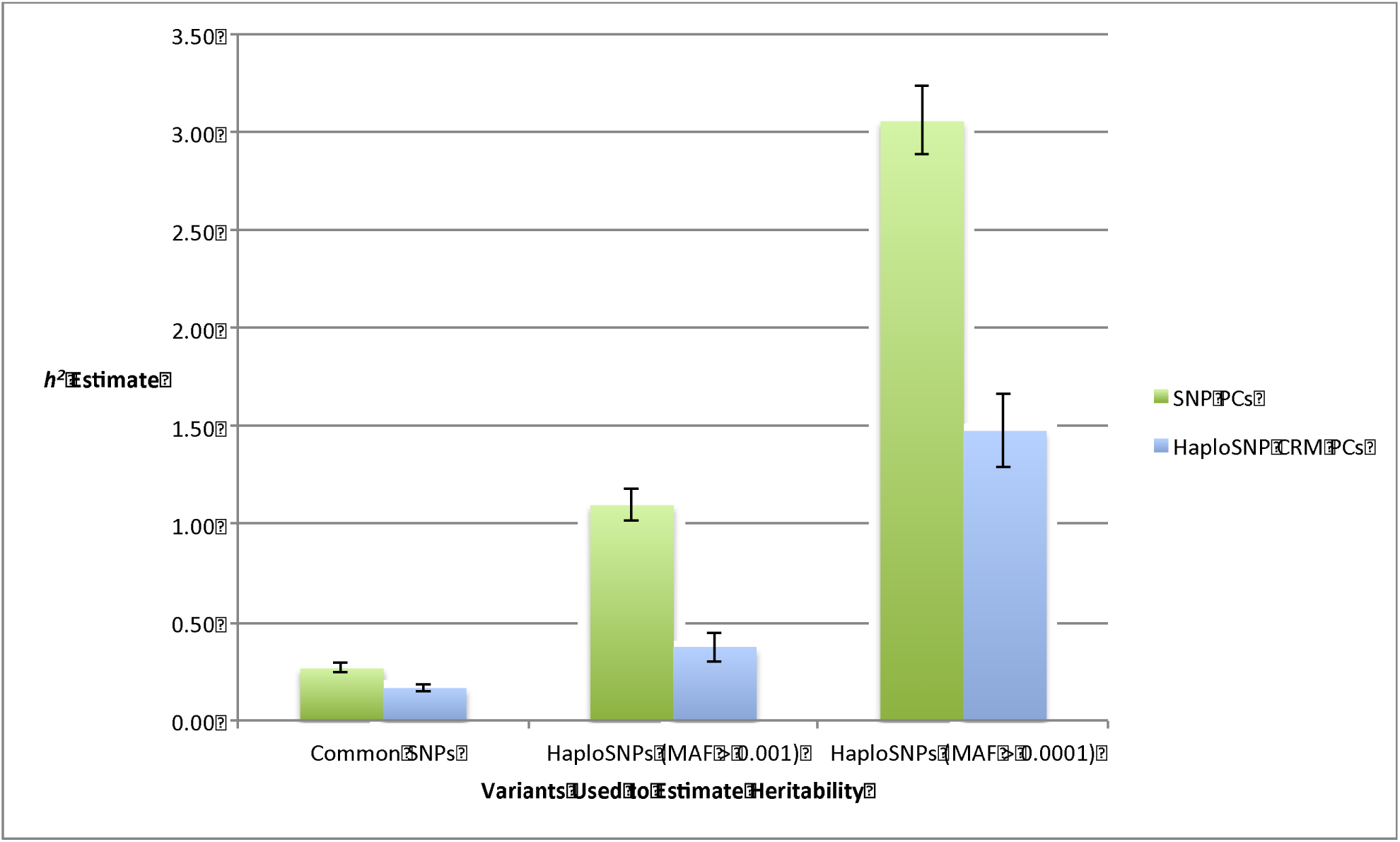
Estimates of heritability of multiple sclerosis explained by common SNPs and haploSNPs in WTCCC2 data. We analyze the heritability of multiple sclerosis explained by array SNPs and haploSNPs (all estimates on the liability scale). Relative to SNP PCs, inclusion of haploSNP PCs produces reductions in heritability estimated from haploSNPs and array SNPs. Standard errors are indicated on the plot. Notably, the estimate of 0.27 (s.e. 0.02) obtained from array SNPs after including PC covariates is 1033 consistent with previous estimates^10,33^, but inclusion of haploSNP PCs reduces this estimate to 0.17 (s.e. 0.02) suggesting that previous estimates may be inflated due to by subtle population stratification. We note that despite the large reductions in 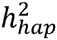 (for both MAF > 0.0001 and MAF > 0.001), it is very likely that these estimates are still inflated by subtle stratification. As a result, we cannot conclude that haploSNPs explain more heritability than genotyped SNPs alone.

### Heritability explained by haploSNPs

We estimated the heritability explained by a subset of the haploSNPs described above with (MAF > 0.001). We obtained an estimate of 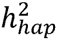 = 1.10 (s. e. 0.08) when including PCs from array SNPs as covariates; this estimate is outside the plausible 0-1 range, suggesting severe confounding. The estimate decreased substantially to 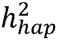 = 0.37 (s.e. 0.07) after correcting for PCs from haploSNP CRMs (along with PCs from array SNPs).

Despite the large reduction in estimates of 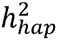 after correcting for PCs from haploSNP CRMs, residual confounding remained a concern. To test for this confounding we expanded our analysis to estimate the heritability explained by all 53.0M haploSNPs (MAF > 10^-4^) described above, and obtained estimates of 3.06 (s. e. 0.18) when correcting for PCs from array SNPs and 1.48 (s. e. 0.19) when correcting for PCs from haploSNP CRMs (along with PCs from array SNPs) (see Figure 2); these values are both outside the plausible 0-1 range, indicative of severe confounding even after correcting for PCs from haploSNP CRMs. This confirms the existence of residual confounding in estimates of heritability explained by haploSNPs with MAF > 10^-4^, and suggests that estimates from more common haploSNPs may continue to be inflated. As a result, we view estimates of heritability explained by haploSNPs in the WTCCC2-MS data set with caution.

## Discussion

By analyzing control-control heritability in the well-studied UK10K data set^8^, we demonstrated that estimates of heritability explained by sequenced SNPs, imputed SNPs and haploSNPs—haplotype variants constructed from within the sample—can be severely inflated. Given that we observed confounding at rare variants not subject to sequencing or imputation error, we believe that subtle stratification, rather than assay artifact or imputation error, is the most likely source of the confounding. This stratification is immune to standard methods of correction: inclusion of principal components (PCs) from array SNPs, imputed SNPs or sequence SNPs. While our results show that PCs from haploSNP CRMs do significantly reduce the impact of confounding, they are unable to control for it entirely. Association statistics at rare variants are also likely to be inflated by this subtle stratification^14^, although heritability estimates aggregate the effects of millions of variants and may be more strongly confounded. UK10K control-control heritability estimates from array SNPs were also inflated, although to a lesser degree, and the extremely subtle nature of the stratification prevented a sum of per-chromosome estimates approach^28^ from detecting confounding. Indeed, this suggests that UK10K control-control stratification is subtler than in previously discussed scenarios^34,35^, where an approach similar to the sum of per-chromosome estimates approach worked well^36^. (We note that a recent paper^37^ raised broader concerns about GCTA^4,24^ and related methods, which, if valid, would render much of our work moot; however, that paper contains 8 errors that invalidate its theoretical and empirical conclusions; see Supplementary Note).

We also observed significant evidence of stratification in our analysis of 22 randomly ascertained phenotypes from the GERA cohort. Notably, if we had used standard methods to correct for this confounding (i.e. including PCs from array SNPs or imputed SNPs), we would have incorrectly concluded that imputed SNPs explained significantly more heritability of the studied traits. This suggests that even in randomly ascertained studies—protected against assay artifact and stratification induced by the ascertainment process—subtle stratification may still confound heritability estimates from imputed SNPs. Estimates of heritability explained by haploSNPs (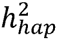) were similarly confounded in the GERA cohort, and showed more extreme evidence of confounding in our analyses of ascertained case-control traits. In our analyses of both schizophrenia and multiple sclerosis, we observed large reductions in 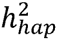 after correcting for PCs from haploSNP CRMs, though uncorrected confounding may continue to inflate these estimates. Correction for PCs from haploSNP CRMs also reduced estimates of 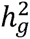 for all three data sets, though to a much lesser degree.

Overall, current methods may be unable to fully quantify or correct for confounding in estimates of heritability from rare variants, although PCs computed from haploSNP CRMs may be able to provide an indication that inflation due to stratification is a concern. Until a lack of inflation due to stratification can be confirmed, we suggest that estimates of heritability from rare variants be viewed with caution. For analyses of array SNPs, we recommend application of LD score regression^18^ to detect confounding, and correction for PCs computed from haploSNP CRMs if subtle stratification is a concern. Despite the potential for uncorrected stratification to inflate estimates of heritability explained by common SNPs, multiple lines of evidence, including enrichment of heritability in biologically relevant parts of the genome^20,38,39^ and strong genetic correlation between studies of the same trait^9^ and across traits^40,41^, suggest that the bulk of estimated 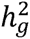 for most studied traits is due to true polygenic signal.

## URLs

HaploSNP Software: http://www.hsph.harvard.edu/faculty/alkes-price/software/Efficient PCGC Regression Software: http://www.hsph.harvard.edu/faculty/alkes-price/software/

## Acknowledgements

We are grateful to Kenneth S. Kendler, Michael C. O'Donovan, Naomi Wray, and Peter Visscher for helpful discussions. This research was funded by NIH grants R01 HG006399 and R01 MH101244.

## Online Methods

### Definition of heritability in the presence of population structure

Following ref. 20 we define heritability explained by a set of variants as the maximum *γ*^2^ between the true phenotype and a linear combination of these variants. However, in the setting of population structure, this definition needs to be modified to include population label as a covariate. Specifically, assuming that values of genetic ancestry *γ*_1_,*γ*_2_ … *γ*_*k*_ for *k* PCs are known, we fit a linear combination of variants and population labels:

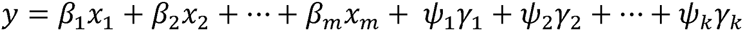

to obtain effect sizes *β*_1_, *β*_2_ … *β*_*m*_ for SNPs and *ψ*_1_, *ψ*_2_ … *ψ*_*m*_ for genetic ancestries. Using these, we define the heritability explained by the *m* variants as:

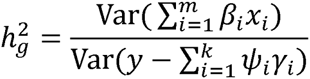

We note that this is a population-level parameter that does not depend on any finite sample. For small values of *F*_ST_ (< 0.01) current methods provide nearly unbiased estimates of this parameter in settings with and without environmental stratification, though estimation becomes biased as *F*_ST_ grows large (see Table S12).

### Estimating heritability with covariates

We used Haseman-Elston (HE) regression^21^ to estimate heritability in control-control analyses. In the standard regression, off-diagonal entries of the phenotypic covariance matrix are regressed against off-diagonal entries of the genetic covariance matrix. The regression coefficient for the GRM is an estimate of the heritability explained by the variants used to construct the GRM. To control for population stratification, we regressed PC covariates out of the phenotype, and included pseudo-GRMs computed from these PCs in the regression. These GRMs were calculated as 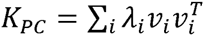 where λ_i_ is the eigenvalue corresponding to eigenvector *λ*_i_. PCs were only combined into the same pseudo-GRM if they were computed from the same set of variants.

For our analyses of case-control phenotypes, we used PCGC regression^22^—a recent generalization of HE regression used to produce estimates on the liability scale accounting for case-control ascertainment. The analyses using PCGC regression were similar to those described above, but covariates were regressed out of the phenotype using logistic regression, and the regression was adjusted to convert estimates to the liability scale as described in ref. 22. Estimates were produced using our previously 22 published efficient implementation of PCGC regression (see URLs), to enable analyses of large data sets^31^. Standard errors for all analyses were computed by jackknifing over individuals^22^.

### MAF-partitioned heritability estimation

In rare variant analyses, we dealt with potential bias in heritability estimates introduced by MAF-dependent genetic architectures^36^ by partitioning variants by MAF and estimating heritability jointly for all MAF bins^42^. We note that MAF partitioning is not robust to LD bias in genetic architecture that is not MAF-mediated, and may not eliminate bias if the genetic architecture is not fully modeled by the MAF bins employed^43^. More complex methods for addressing LD bias have been proposed^10,36^, though the suitability of these methods for analyses of rare variants is unclear^11^. While possible uncorrected LD bias is a concern, we note that this bias would affect estimates with and without covariates and would be unlikely to alter our conclusions.

MAF bins were chosen based on the data set analyzed, as sample sizes varied. We computed PCS from 7 MAF bins of sequenced SNPs, imputed SNPs, and haploSNPs in the UK10K data: [doubleton-0.0005], (0.0005-0.001],(0.001-0.01], (0.01-0.05], (0.05-0.1], (0.1-0.25], and (0.25-0.5]. For heritability estimation in this data set, we used the 5 MAF bins with MAF > 0.001. When analyzing haploSNPs and imputed SNPs in the PGC2-SCZ, WTCCC2-MS, and GERA data sets, we considered MAF bins of [0.0001-0.001], (0.001-0.005], (0.005-0.01], (0.01-0.05], (0.05-0.1], (0.1-0.25], and (0.25-0.5]. For heritability estimation in the WTCCC2-MS and GERA data sets we used all 7 MAF bin; for the PGC2-SCZ data we used only the 6 MAF bins with MAF > 0.001. We note that none of the individual cohorts in the PGC2-SCZ data had > 10,000 individuals, so the lowest MAF bin (used only for PC computation) included all non-singleton variants. All analyses with array SNPs used only a single variance component.

### Constructing haploSNPs

HaploSNPs are haplotypes of adjacent SNPs excluding a subset of masked sites that arise from skipped mismatches. Individuals are defined to carry 0, 1, or 2 copies of the haploSNP if none, one or both of their chromosomes matches the haplotype at all unmasked sites. The algorithm to generate haploSNPs (see Supplementary Note for pseudocode) proceeds from phased genotypes. For our haploSNP analysis, we used the HAPI-UR method^25^ to computationally phase genotypes. Using these phased genotypes we build a set of haplotype variants. At each polymorphic SNP, we create two haploSNPs—one for the ancestral allele and one for the derived allele. We expand these haploSNPs until a terminating mismatch is detected. A terminating mismatch is one that cannot be explained without a recombination between the current haploSNP and the mismatch SNP. This is tested using a standard 4-gamete test^44^. Once a terminating mismatch is detected, we terminate the current haploSNP and create two child haploSNPs: one for individuals that match the current haploSNP and the ancestral allele at the mismatch SNP, the other for individuals that match the current haploSNP and the derived allele at the mismatch SNP. We repeat this process until the current haploSNP is longer than a length threshold, or has MAF lower than a MAF threshold.

The output of this algorithm is a list of haploSNPs. These haploSNPs are a mapping of multiple co-located SNPs to a particular allele at each SNP. For each haploSNP, each phased chromosome is assigned either a 1, indicating a perfect match at all SNPs that make up the haploSNP, or a 0 otherwise. This set of biallelic haploSNPs is then used in downstream analysis in addition to biallelic SNPs. We note that all quality control steps (see below) are applied to SNPs prior to construction of haploSNPs, no additional QC steps are applied to the haploSNPs in our analysis.

We note that prior work on haplotype association analyses^45-48^ has focused on analyses of a small number of co-located SNPs (<10) for the purposes of identifying associations between combinations of these SNPs and phenotypes. While these approaches are substantially different than our method for generating haploSNPs, association statistics for rare haplotypes produced by these methods may be vulnerable to the effects of subtle stratification that we observe here.

### Constructing correlation relationship matrices (CRM)

Correlation matrices are used to compute rare variant PCs because covariance matrices may have large diagonal entries that result in outlier PCs. Entries of the standard GRM from a set of variants *S* are computed as:

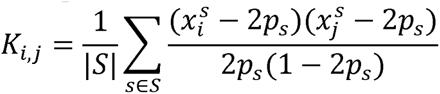

We compute the correlation relationship matrix by normalizing the standard GRM by the appropriate diagonal entries

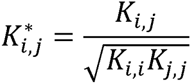

### Imputation of rare variants

To perform imputation, we computationally phased our genotypes^25^, and used the Impute2 software package^29^ to produce a set of imputed variants. While it is common practice to restrict analyses to well-imputed (e.g. INFO > 0.9) genotypes, a recent analysis^11^ suggests that poorly imputed SNPs may better tag unobserved genotypes and explain substantially more heritability than genotyped SNPs alone. To use these variants, we convert the genotype probabilities output by Impute2 into hard genotype calls by calling the max-likelihood genotype as ground truth. Specifically we use the plink2 software package^49^ with the command line option: ‐‐hard-call-threshold 0.4999.

### Data sets

#### UK10K data set

The UK10K project^8^ data is comprised of low-coverage sequencing (7x) from individuals from two cohorts: ALSPAC and TWINSUK. We combined these data sets and applied stringent QC, removing SNPs that had either a deviation from Hardy-Weinberg equilibrium at a p-value below 0.01, or missingness greater than 0.002. For our analyses of sequenced SNPs we did not impose any threshold on minor allele frequency, and were left with 17.6M non-singleton sequenced SNPs. We used the full set of sequenced SNPs to compute PCs for controlling population stratification and 11.7M sequenced SNPs with MAF > 0.001 to estimate heritability from sequenced SNPs. We then focused on the set of SNPs that were also typed on the Illumina Human660-Quad chip genotyping array (used by WTCCC2^23^). These SNPs were defined “array SNPs” in our analyses. We also removed one individual in any pair of individuals with relatedness greater than 0.025 by array SNP covariance. Following all QC steps, we analyzed 3,565 individuals—1,817 from ALSPAC and 1,748 from TWINSUK, using 408k array SNPs. We computationally phased these genotypes^25^, and used the phased genotypes to impute 17.4M non-singleton SNPs of which 13.0M had MAF > 0.001. We also used these phased genotypes to construct a total of 32.3M haploSNPs of which 26.5M had MAF > 0.001.

#### GERA data set

The GERA data set is comprised of genotype data from the GERA EUR chip and phenotype data for each of 22 disease conditions based on electronic medical records for 54,734 individuals of European ancestry^31^. We averaged heritability estimates of 22 randomly ascertained case-control phenotypes recorded as part of the GERA data set^19^ (see Table S8). While we expect assay artifact to be largely uncorrelated with phenotype in randomly ascertained case control-studies, we used stringent QC to ensure that it did not impact our estimates. Specifically, we removed any SNPs that had either a deviation from Hardy-Weinberg equilibrium at a p-value below 0.01, or missingness greater than 0.002. We also removed one individual in any pair of individuals with relatedness greater than 0.025 by SNP covariance. Following all QC steps, we analyzed 47,360 individuals 25 genotyped at 289k SNPs. We computationally phased these genotyped SNPs^25^ and used these phased genotypes to impute (see below) 22.6M SNPs of which 13.7M had MAF > 0.001. We also built a set of 39.3M haploSNPs (see below) of which 27.5M had MAF > 0.001.

#### Schizophrenia data set (PGC2 data set)

The PGC2^32^ data set is comprised of several cohorts of diverse ancestry that were genotyped on a variety of different genotyping platforms. To avoid issues related to cross-population heritability estimation, we focused on meta-analysis of estimates within cohorts. Our estimates were produced from each of 10 cohorts with >1,000 individuals. Within each cohort we applied stringent QC to genotyped SNPs (we did not analyze imputed SNPs), removing any SNPs that were below 0.01 minor allele frequency, had deviation from Hardy-Weinberg equilibrium at a p-value below 0.01, had missingness greater than 0.002, or had differential missingness between cases and controls with a *p*-value below 0.05. We also removed one individual in any pair of individuals with relatedness greater than 0.025 by SNP covariance. Following all QC steps, we analyzed 35,238 individuals. This is smaller than the number of individuals analyzed the largest previous meta-analysis^32^ because we restricted to individuals that came from cohorts with >1,000 individuals. The average number of SNPs genotyped in each cohort was 461k. We computationally phased these genotyped SNPs^25^ and built an average of 32.8M haploSNPs in each cohort (see Table S10), of which an average of 26.3M had MAF > 0.001.

#### Multiple sclerosis data set (WTCCC2 data set)

We analyzed the publicly available subset^26^ of data analyzed in a large GWAS of multiple sclerosis^23^. As cases and controls were genotyped separately, we used a very high level of stringency in our quality control. Specifically, we removed any SNPs that had minor allele frequency below 0.02, had deviation from Hardy-Weinberg equilibrium at a p-value below 0.05, had missingness greater than 0.002, or had differential missingness between cases and controls with a p-value below 0.05. We also removed one individual in any pair of individuals with relatedness greater than 0.05 by SNP covariance. We subsequently performed five rounds of outlier removal whereby all individuals more than 6 standard deviations away from the mean along any of the top 20 eigenvectors were removed and all eigenvectors recomputed. Following all QC steps, we analyzed 14,526 individuals genotyped at 375k SNPs. These individuals consisted of 9,315 cases, 2,635 controls from the NBS cohort and 2,794 controls from the 58C cohort. We computationally phased the genotypes^25^ and built a set of 53.0M haploSNPs (see below) with MAF > 0.0001, of which 36.3M had MAF > 0.001. To avoid biases due to the large effect of the well-known HLA locus, we excluded chromosome 6 from all heritability analyses, leaving a total of 349k SNPs. While the effect of the HLA locus is the largest in the genome, it has been estimated to explain only about 3% of the phenotypic variance of MS on the liability scale^33^, and thus should not affect our results substantially.

